# Appropriate Correction for Multiple Comparisons in Decoding of ERP Data: A Re-Analysis of Bae & Luck (2018)

**DOI:** 10.1101/672741

**Authors:** Gi-Yeul Bae, Steven J. Luck

## Abstract

Bae & Luck (2018) reported a study of visual working memory in which the orientation being held in memory was decoded from the scalp distribution of sustained ERP activity and alpha-band EEG oscillations. Decoding accuracy was compared to chance at each point during the delay interval, and a correction for multiple comparisons was applied to find clusters of consecutive above-chance time points that were stronger than would be expected by chance. However, the correction used in that study did not account for the autocorrelation of the noise and may have been overly liberal. Here, we describe a more appropriate correction procedure and apply it to the data from Bae & Luck (2018). We find that the major clusters of time points that were significantly above chance with the original correction procedure remained above chance with the updated correction procedure. However, some minor clusters that were significant with the original procedure were no longer significant with the updated procedure. We recommend that future studies use the updated correction procedure.

---

A well-known advantage of EEG-based measures of neural activity over other noninvasive measures is their millisecond-level temporal resolution. However, this advantage brings with it the need to assess statistical significance at each of many time points, which can lead to an inflation of Type I error rates if an appropriate correction for multiple comparisons is not applied. An increasingly common approach is to search for clusters of consecutive time points that are individually significant without correction and determine whether the cluster as a whole is larger than would be expected by chance (Groppe, Urbach, & Kutas, 2011b, 2011a; Maris & Oostenveld, 2007). The cluster size that is beyond what would be expected by chance is typically determined by permuting the labels on the data (to simulate the null hypothesis) a large number of times and determining the null distribution of cluster sizes from these permutations.

Bae and Luck (2018) used a similar approach in a study of visual working memory. In this study, subjects stored the orientation of a teardrop shape in working memory and then reproduced the remembered orientation at the end of the trial. Sixteen different orientations were possible. An SVM-based classification algorithm was then applied to either the averaged ERPs or the averaged alpha-band activity from each orientation to determine whether the orientation of the stimulus could be predicted from the scalp distribution of the neural signal at a given moment in time. Decoding accuracy was above chance for most of the retention interval in most of the analyses. Clusters of significant time points were determined using a Monte Carlo approach that made a random guess about the orientation of the stimulus for each of many iterations, and this was then used to determine the largest cluster of individually significant time points that would be expected by chance.

However, the Monte Carlo approach treated each time point as an independent sample and did not account for the autocorrelation of the noise in EEG data. For example, if one point has a large voltage, the next point will also likely have a large voltage. Similarly, if the pattern of noise across the scalp at one point differentiates between two orientations, then the pattern of noise across the scalp at the next point will also likely differentiation between these two orientations. This will tend to increase the likelihood of obtaining multiple consecutive time points that have high decoding accuracy (as well as multiple consecutive time points that have low decoding accuracy). If the model of the null hypothesis does not take this into account, then it will underestimate the cluster sizes that will be produced by chance, making the statistical test too liberal.

The goal of the present report was to describe a more appropriate cluster-based correction for multiple comparisons that accounts for the temporal autocorrelation and apply this improved correction to the data of Bae and Luck (2018).

## Method

The key to taking noise autocorrelation into account is to use permutations of the actual data to construct a null distribution, making sure to permute all time points for a given EEG epoch together so that the autocorrelated noise is present in the permuted data. This permutation approach was taken by similar studies of working memory, such as Foster et al. (2016). The goal is to construct a null distribution that reflects what would be expected if all the trials (e.g., for all the different orientations) were actually sampled from a single distribution. Randomly permuting the labels before computing the test statistic achieves this. In some approaches, the permutation is performed prior to training the classifier. However, the training stage can be quite time-consuming, and performing a large number (e.g., 1000) permutations at this stage can be prohibitively slow.

Our updated method therefore involves performing a permutation at the time of test. That is, after the classifier has been trained on the non-permuted data, the labels on the test data are permuted prior to determining classification accuracy. With this approach, the classifier is necessarily guessing randomly, making it possible to construct a null distribution by performing a large number of permutations. Crucially, because the same permuted label was assigned for each time point in a given EEG epoch, any temporal autocorrelation in the data was present during the construction of the estimated null distribution. We have used this method in two other studies (Bae & Luck, in press, 2019), and we refer the reader to Bae & Luck (2019) for a detailed description of the decoding method and the statistical analysis method. For the present article, we simply applied this updated statistical method to the data of Bae and Luck (2018).

## Results

Figure 1 shows decoding accuracy at each point in time following the sample stimulus in Experiment 1 of Bae and Luck (2018). On each trial of this experiment, subjects were shown a 200-ms teardrop in one of 16 orientations, maintained the orientation of the teardrop in memory for 1300 ms, and then attempted to reproduce the orientation of the teardrop. An SVM-based multiclass decoding algorithm was used to decode the orientation of the stimuli on the basis of the scalp distribution at a given time point. Decoding was done separately for the sustained ERP activity and for the alpha-band EEG activity. A three-fold cross-validation procedure with multiple iterations was used to prevent overfitting.

**Figure 1.**
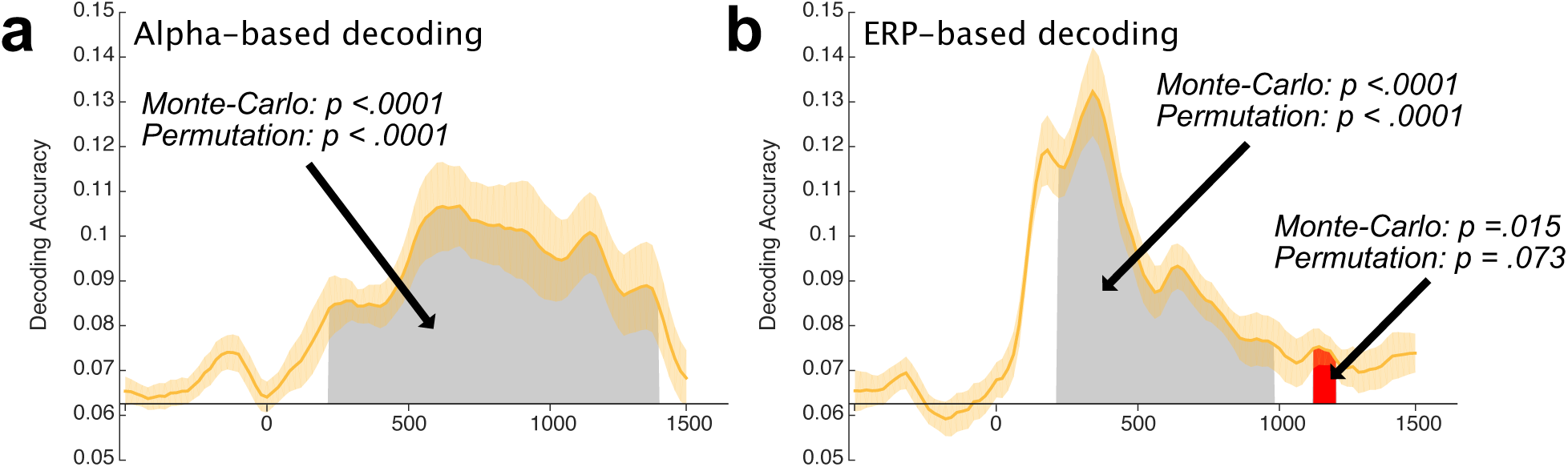
Mean accuracy of (a) alpha-based decoding and (b) ERP-based decoding in Experiment 1. Chance-level performance (1/16 = .0625) is indicated by the X axis line. The orange shading indicates ±1 SEM. Gray areas indicate clusters of time points in which the decoding was significantly greater than chance after correction for multiple comparisons using both the Monte Carlo method and the permutation method. Red areas indicate clusters of time points in which the decoding was significantly greater than chance after correction for multiple comparisons using the Monte Carlo method but not the permutation method.

Decoding accuracy was compared to chance (1/16) at each time point between the offset of the sample stimulus (200 ms) and the end of the retention interval (1500 ms). Gray areas in the figure show clusters of points that were statistically significant as determined both by the original Monte Carlo statistical method and the updated permutation method, and red regions show clusters of points that were significant with the original method but not with the updated method. The updated method is more conservative, so there were no clusters that were significant with the updated method but not with the original method. The *p* value for each cluster is also shown in the figure. For both alpha-based and ERP-based decoding, a large cluster of points that spanned most of the retention interval was significant with both the original and the updated methods. The only difference was a small cluster of points near the end of the retention interval in the ERP-based decoding that was significant with the original Monte Carlo method (*p* = .015) but was not significant (*p* = .073) with the updated permutation method.

Figure 2 shows decoding accuracy at each point in time following the sample stimulus in Experiment 2 of Bae and Luck (2018). In this experiment, the sample stimulus was one of 16 different orientations that was presented in one of 16 different locations; orientation and location were independently chosen on each trial. The task required reproducing the orientation of the stimulus at a randomly-chosen location. Thus, location was completely task-irrelevant. We independently decoded the location of the stimulus (collapsed across orientations) and the orientation of the stimulus (collapsed across locations).

**Figure 2.**
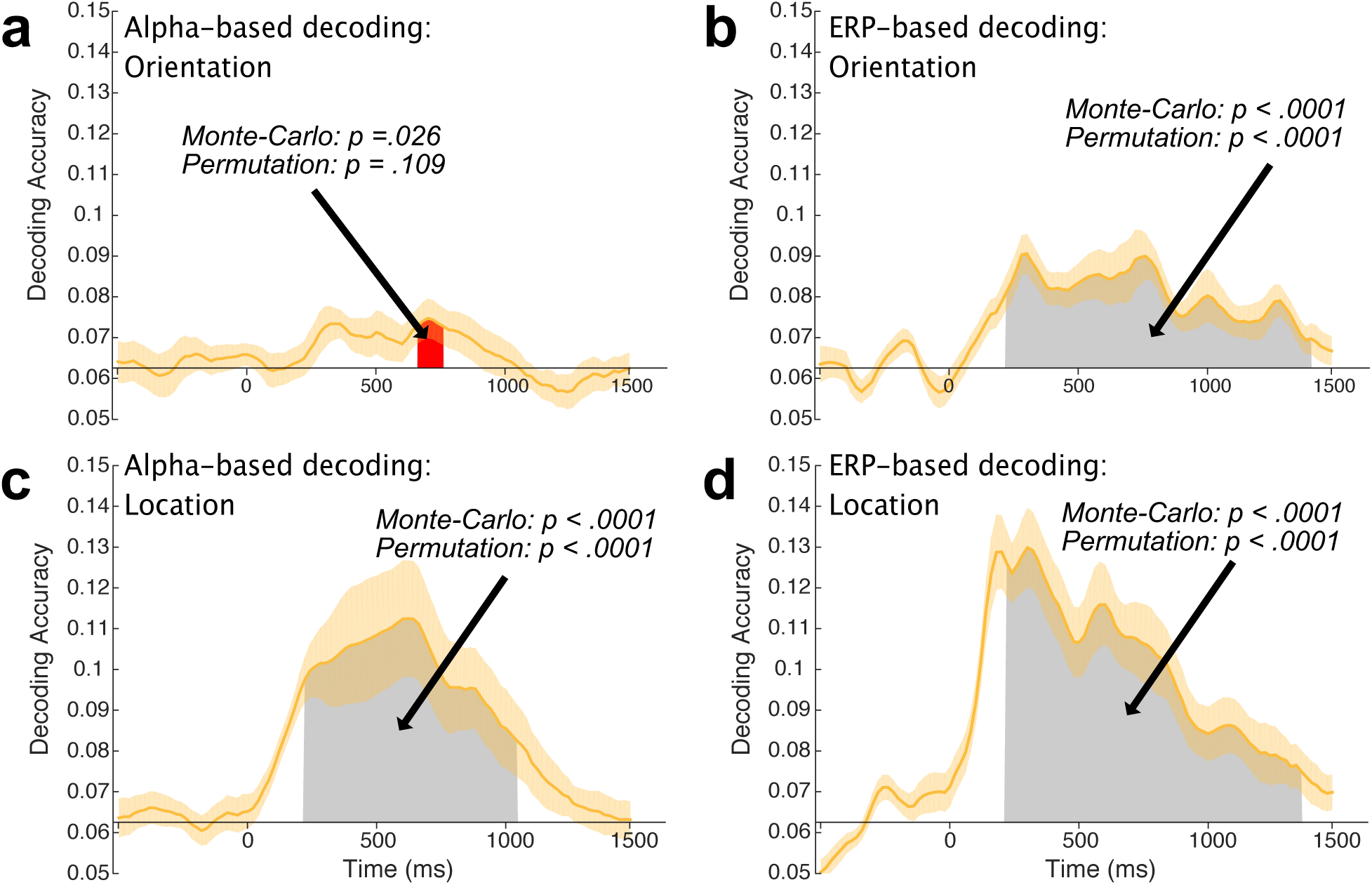
Alpha-based decoding accuracy for (a) the orientation of the sample teardrop and (c) the location of the sample teardrop tip. ERP-based decoding accuracy for (b) the orientation of the sample teardrop and (d) the location of the sample teardrop tip. The orange shading indicates ±1 SEM. Gray areas indicate clusters of time points in which the decoding was significantly greater than chance after correction for multiple comparisons using both the Monte Carlo method and the permutation method. Red areas indicate clusters of time points in which the decoding was significantly greater than chance after correction for multiple comparisons using the Monte Carlo method but not the permutation method.

As shown in Figure 2, the pattern of significance was nearly identical for the original Monte Carlo statistical method and the updated permutation method. For the ERP-based decoding of both location and orientation, a large cluster of points was significantly greater than chance irrespective of the statistical method. Alpha-based decoding of location produced a large cluster of points with both statistical methods. For alpha-based decoding of orientation, decoding accuracy was quite low throughout the retention interval, with a small cluster that was significant (*p* = .026) with the original statistical method but not with the updated method (*p* = .109).

In Experiment 2, we also asked if we could decode the orientation of the stimulus by training on the data from one set of locations and testing on the data from another set of locations. We similarly asked if we could decode the location of the stimulus by training on the data from one set of orientations and testing on the data from another set of orientations. The results of these cross-feature decoding analyses are shown in Figure 3. No clusters of significant decoding accuracy were observed using either statistical method when we attempted to decode the orientation of the stimulus from the alpha-band activity. For the other three cases, both statistical methods found clusters of significantly above-chance decoding. However, a few additional clusters were significant for the original method but not for the updated method.

**Figure 3.**
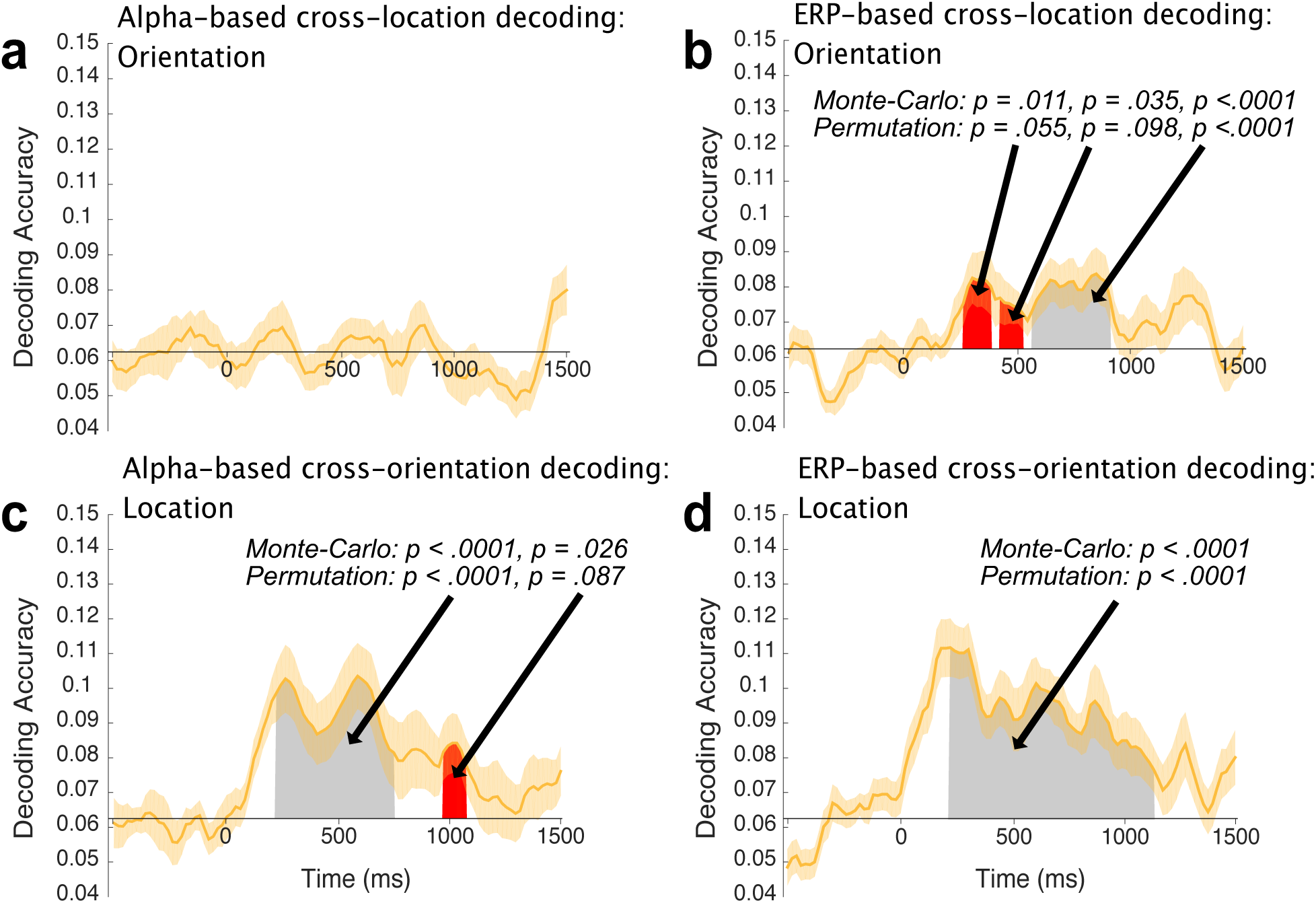
Average cross-feature decoding accuracy at each time point. (a) Average accuracy of alpha-based cross-location decoding of orientation. (b) Average accuracy of ERP-based cross-location decoding of orientation. (c) Average accuracy of alpha-based cross-orientation decoding of location. (d) Average accuracy of ERP-based cross-orientation decoding of location. The orange shading indicates ±1 SEM. Gray areas indicate clusters of time points in which the decoding was significantly greater than chance after correction for multiple comparisons using both the Monte Carlo method and the permutation method. Red areas indicate clusters of time points in which the decoding was significantly greater than chance after correction for multiple comparisons using the Monte Carlo method but not the permutation method.

## Discussion

These results show the importance of considering the autocorrelation of the noise when correcting for multiple comparisons in assessing consecutive time points with EEG data. If the autocorrelation is not taken into account, the statistical method is overly liberal.

Fortunately, the application of the more appropriate permutation-based test to the data of Bae and Luck (2018) indicated that the main conclusions of that study were sound. Although a few small clusters of points were no longer statistically significant when the permutation-based approach was applied, this updated method yielded at least one major cluster of significant decoding in each condition where the original method yielded significance.

## References

Bae, G. Y., & Luck, S. J. (in press). Reactivation of previous experiences in a working memory task. Psychological Science.

Bae, G. Y., & Luck, S. J. (2018). Dissociable Decoding of Working Memory and Spatial Attention from EEG Oscillations and Sustained Potentials. Journal of Neuroscience, 38, 409–422.

Bae, G. Y., & Luck, S. J. (2019). Decoding motion direction using the topography of sustained ERPs and alpha oscillations. NeuroImage, 184, 242–255.

Foster, J. J., Sutterer, D. W., Serences, J. T., Vogel, E. K., & Awh, E. (2016). The topography of alpha-band activity tracks the content of spatial working memory. Journal of Neurophysiology, 115, 168–177.

Groppe, D. M., Urbach, T. P., & Kutas, M. (2011a). Mass univariate analysis of event-related brain potentials/fields I: a critical tutorial review. Psychophysiology, 48, 1711–1725.

Groppe, D. M., Urbach, T. P., & Kutas, M. (2011b). Mass univariate analysis of event-related brain potentials/fields II: Simulation studies. Psychophysiology, 48, 1726–1737.

Maris, E., & Oostenveld, R. (2007). Nonparametric statistical testing of EEG-and MEG-data. Journal of Neuroscience Methods, 164, 177–190.

